# Integrating thermodynamic and enzymatic constraints into genome-scale metabolic models

**DOI:** 10.1101/2020.11.30.403519

**Authors:** Xue Yang, Zhitao Mao, Xin Zhao, Ruoyu Wang, Peiji Zhang, Jingyi Cai, Hongwu Ma

## Abstract

Stoichiometric genome-scale metabolic network models (GEMs) have been widely used to predict metabolic phenotypes. In addition to stoichiometric ratios, other constraints such as enzyme availability and thermodynamic feasibility can also limit the phenotype solution space. Extended GEM models considering either enzymatic or thermodynamic constraints have been shown to improve prediction accuracy. In this paper, we propose a novel method that integrates both enzymatic and thermodynamic constraints in a single Pyomo modeling framework (ETGEMs). We applied this method to construct the EcoETM, the *E. coli* metabolic model iML1515 with enzymatic and thermodynamic constraints. Using this model, we calculated the optimal pathways for cellular growth and the production of 22 metabolites. When comparing the results with those of iML1515 and models with one of the two constraints, we observed that many thermodynamically unfavorable and/or high enzyme cost pathways were excluded from EcoETM. For example, the synthesis pathway of carbamoyl-phosphate (Cbp) from iML1515 is both thermodynamically unfavorable and enzymatically costly. After introducing the new constraints, the production pathways and yields of several Cbp-derived products (e.g. L-arginine, orotate) calculated using EcoETM were more realistic. The results of this study demonstrate the great application potential of metabolic models with multiple constraints for pathway analysis and phenotype predication.

## 1. Introduction

Constraint-based metabolic network modeling is a mathematical framework used to analyze the feasible metabolic flux solution space through constrained optimization methods (Bordbar et al., 2014). It has been widely used in genome-scale metabolic network analysis to calculate the optimal synthesis pathways, as well as predict growth phenotypes and modification targets for metabolic engineering or disease treatment (Kim et al., 2012). Initially, only stoichiometric constraints and reaction reversibility constraints were considered in a classical method called flux balance analysis (FBA) (Orth et al., 2010). With the accumulation of enzyme kinetics data and the availability of high-throughput omics data, it has become possible to incorporate these data into the models to add boundary constraints for individual reactions or a summarized constraint of enzyme resources (Liu et al., 2014). In 2007, the FBAwMC model was constructed by introducing constraints of enzyme resources based on a fixed cell volume (Beg et al., 2007). Subsequently, other integration methods of protein resources were developed (Yang et al., 2018). There are two major trends in the development of resource allocation models. One is the MOMENT (Adadi et al., 2012) type models with only enzymatic constraints on the basis of GEMs, while the other is ME (Lloyd et al., 2018) type models with more detailed description of cellular processes, such as transcription and translation. In 2017, Sanchez et al. reported GECKO (GEMs with Enzymatic Constraints using Kinetic and Omics data) method and applied it in the construction of an enzymatic constraints model of *Saccharomyces cerevisiae* (Sanchez et al., 2017). This method was soon extended and applied in the construction of enzymatic constraints GEMs (ECGEMs) of other species (Bekiaris and Klamt, 2020; Massaiu et al., 2019). By integrating *k*_cat_ parameters for individual enzymes and total enzyme amount constraints, these models can improve the simulation and prediction of biological phenomena, such as overflow metabolism (Molenaar et al., 2009) and pathways switching (Chen and Nielsen, 2019).

In FBA models, certain reactions are set as irreversible by considering the thermodynamic feasibility by introducing a zero-value constraint on upper/lower bounds of a reaction. However, there are no clear criteria to determine whether a reaction should be reversible or not, and reactions that are thermodynamically feasible by themselves can form thermodynamically unfavorable pathways such as unlimited ATP generation loops (Yuan et al., 2017). To address this problem, methods combining thermodynamic constraints with GEMs have been developed to improve the prediction accuracy (Soh and Hatzimanikatis, 2010). In 2007, Henry et al. integrated thermodynamic constraints into the FBA calculation process and proposed the TFMA method (Henry et al., 2007). Recently, Salvy et al. developed this method into the pyTFA and matTFA toolkits (Salvy et al., 2019) and applied it to phenotypic analysis in combination with the ME model (Salvy and Hatzimanikatis, 2020). Reliable data on thermodynamic parameters of reactions is particularly important for models with thermodynamic constraints (Du et al., 2018; Noor et al., 2012). In 2011, Flamholz et al. developed the eQuilibrator, a biological thermodynamics calculator that enables users to easily obtain thermodynamic parameters (Flamholz et al., 2012). In 2014, Noor et al. introduced the concept of Max-min Driving Force (MDF) to predict and optimize the thermodynamic bottleneck reactions in a pathway, and integrated this function into the eQuilibrator website as a free tool (Noor et al., 2014). Based on these studies, Hadicke et al. proposed the optMDFpathway method, which can directly identify the optimal MDF (and hence the most thermodynamically feasible) pathways in GEMs (Hadicke et al., 2018). Different from some workflows such as Poppy (Asplund-Samuelsson et al., 2018), which requires defining the pathway in advance and then evaluating its thermodynamic driving force, the optMDFpathway method integrates the objects of MDF into the FBA solution process and can therefore be directly applied to GEMs.

In this paper, we propose a novel method that integrates both enzymatic and thermodynamic constraints into a single modeling framework, named ETGEMs. The Python-based Pyomo modeling package (Hart et al., 2017; Hart et al., 2011) was used to integrate multiple objects and constraints to satisfy the different expectations of the optimal pathways, such as maximal yield, minimal enzyme cost and optimal thermodynamic driving force. We applied this method to construct EcoETM, an *E. coli* metabolic model with enzymatic and thermodynamic constraints based on the iML1515 model (Monk et al., 2017). The simulation results indicated that the new model can effectively reduce the solution space by excluding pathways that are thermodynamically unfavorable or have high enzyme costs exceeding the available resources. The integration of both thermodynamic and enzymatic constraints into a genome-scale metabolic network model, the ETGEMs modeling framework, can be applied to other organisms with available enzyme kinetics and reaction thermodynamics data. The code for the construction and analysis of the model is available at https://github.com/tibbdc/ETGEMs.

## 2. Methods

### 2.1. Pretreatment of the initial model and data collection

The *E.coli* iML1515 (Monk et al., 2017) model was selected as the initial model for the integration of constraints and the range of reactions set for the collection of kinetic parameters. All model construction and analysis was conducted using Python (version 3.6.5). The “convert_to_irreversible” function in the Cobrapy toolkit (version 0.13.1) was used to split the reversible reactions, and an irre_iML1515 model was formed. The newly divided reactions were named “original reaction ID_reverse”. The final model contained 3375 one-way reactions, 663 of which were designated as “_reverse”.

Collection of enzymatic parameters: The *k*_cat_ parameters are based on machine learning predictions from databases performed by Heckmann et al. (Heckmann et al., 2018). Among them, a small number of parameters were corrected in previous work according to biomass and product synthesis. The protein subunit composition and molecular weight data were downloaded from the EcoCyc database (https://ecocyc.org/) (Karp et al., 2018). The value of total enzyme amount (e_pool), 0.228 g/gDW, was calculated based on protein abundance data in the PAXdb database (https://pax-db.org/) (Wang et al., 2012) and intracellular protein content of g protein/gDW (Bremer H and P, 1996). An average enzyme saturation value (*σ*) of 0.5 was used based on previous studies (Bennett et al., 2009; Sanchez et al., 2017). The calculation of the enzymatic parameters was reported in a separate paper in detail (https://github.com/tibbdc/ECMpy).

Collection of thermodynamic parameters: the biomass synthesis reactions (2) and transport reactions (Hadicke et al., 2018) (1420) and exchange reactions (361) were excluded first. Among the remaining 1592 reactions, we temporarily removed the 253 “_reverse” reactions. Therefore, the collection range of thermodynamic parameters was reduced to 1339 reactions. The Gibbs energies of reactions were downloaded from the eQuilibrator website (http://equilibrator.weizmann.ac.il/download). After matching KEGG (used in eQuilibrator) and BIGG (used in iML1515) reaction IDs and reaction directions, 586 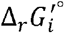 parameters were determined. Then, by referring to Table S5 in previous research (Hadicke et al., 2018), another 145 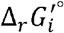 parameters were added. In addition, 71 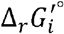 parameters were calculated using the eQuilibrator calculator after manually matching KEGG reaction IDs by unifying reaction equations (e.g. GLCS1: replacing “ADPglucose <=> ADP + Glycogen” with “ADPglucose + 0.25 H2O <=> ADP + 0.25 Glycogen”) and metabolite names (e.g. MLTP1: replacing “Maltopentaose” with “Cellopentaose”). Besides, 123 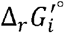 parameters were estimated by referring to similar reactions that can be identified by the eQuilibrator calculator. Among the resulting 925 reactions, 232 reactions had corresponding “_inverse” reactions, and we assigned the 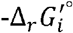 values to their “_inverse” reactions. Finally, a total of 1157 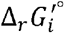 values were obtained, and 435 reactions still lacked 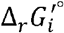 parameters. All 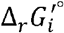 parameters are listed in Tables A-C (in Supplementary file2), and supplementary methods (in Supplementary file1). For the 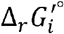 calculated using eQuilibrator, the ionic strength was set to 0.1 M and the pH was set to 7.5. The gas constant *R* was 8.31446 J mol^-1^K^-1^ (Flamholz et al., 2012) and the temperature T was 310.15 K (37 □), giving an RT value of 2.579 kJ/mol.

### 2.2. Setting the concentration range of metabolites

The concentration limits for all metabolites were set to 0.5 μM as lower bound and 20 mM as upper bound (Bennett et al., 2009). The concentrations of CO2 and O2 were more strictly bounded to be in the ranges from 0.1 – 100 μM (Hadicke et al., 2018) and 0.5 – 200 μM (Baltazar Reynafarje et al., 1985; Murphy, 2009), respectively. The concentration ratios for ATP:ADP, ADP:AMP, NAD:NADH, NADPH:NADP and HCO_3_:CO_2_, were respectively fixed to 10:1, 1:1, 10:1, 10:1 and 2:1, based on the literature (Hadicke et al., 2018).

### 2.3. The principle of introducing constraints

Method for stoichiometric and flux balance constraint addition:

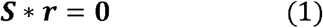

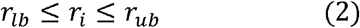

where ***r*** is the reaction flux, and ***S*** represents the stoichiometric matrix (Orth et al., 2010).

A concise method for enzymatic constraint addition (Bekiaris and Klamt, 2020):

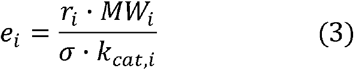

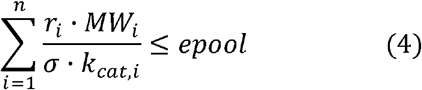

where *e_i_* is the enzyme cost of a reaction flux *r_i_ MW_i_* is the molecular weight of enzyme *i*, and *σ* represents the average saturation of all enzymes.

Method for thermodynamic constraint introduction:

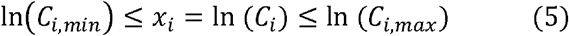

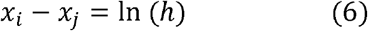

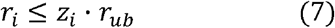

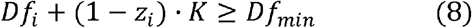

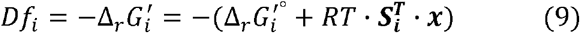

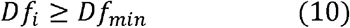

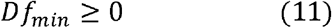

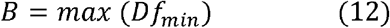

where *h* is the concentration ratio of metabolites *C_i_* and *C_j_*, 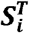 is the transposed *i*-th reaction of the full stoichiometric matrix ***S***. In order to realize the thermodynamic constraints only for the reactions involved in the pathway (*r_i_* > 0), a binary variable *z_i_* and a sufficiently large value *K* (Henry et al., 2007) must be introduced. In this work, the value of *K* was defined as max(*Df_i,max_*) — min(*Df_i,min_*). Due to the second law of thermodynamics, a pathway can only work if formula (11) is valid, When calculating the maximal thermodynamic driving force for implementing the MDF or optMDFpathway methods, it is necessary to set the lower bound of the driving force *Df_i_* as *B* and turn it into an objective function.

### 2.4. Objective functions used in this work

Multiple objective functions were adopted in this work to calculate the optimal pathways satisfying different constraints, as listed in Table 1. In addition, other objective functions were also used for other analyses based on the constrained model, such as calculating the variability of metabolite and enzyme concentrations to identify the bottlenecks in the network. These objective functions are listed in Table 1 and Table S1 (in Supplementary file1).

**Table 1.** Major objective functions used in the modeling framework.

| Objects | Types | Constraints | Purposes |
| --- | --- | --- | --- |
| $r_{biomass}$ or $r_{product}$ | Maximize | $r_{substrate, ub}, epool, x_{i, lb}, x_{i, ub}, h_i, Df_{min}$ | To solve the maximum synthesis rate $r_i$ of biomass or product. |
| $x_i$ | Maximize and Minimize | $r_{substrate, ub}, r_{product, lb}, epool, x_{i, lb}, x_{i, ub}, h_i, Df_{min}$ | To calculate the variability of $C_i$ to find the limiting metabolite. |
| $Df_i$ | Maximize and Minimize | $r_{substrate, ub}, r_{product, lb}, epool, x_{i, lb}, x_{i, ub}, h_i, Df_{min}$ | To calculate the variability of $Df_i$ to find the bottleneck reaction. |
| $(r_i MW_i)/(\sigma k_{cat,i})$ | Maximize and Minimize | $r_{substrate, ub}, r_{product, lb}, epool, x_{i, lb}, x_{i, ub}, h_i, Df_{min}$ | To calculate the variability of enzyme usage to detect key enzymes. |
| $sum [(r_i MW_i)/(\sigma k_{cat,i})]$ | Minimize | $r_{substrate, ub}, r_{product, lb}, epool, x_{i, lb}, x_{i, ub}, h_i, Df_{min}$ | To calculate the minimum enzyme cost of pathways |
| $B$ | Maximize | $r_{substrate, ub}, r_{product, lb},$<br>$epool, x_i, lb, x_i, ub, h_i$ | To calculate the MDF of pathways |

### 2.5. Tools for model construction and problem solving

The Concrete model framework in the python-based Pyomo package (version 5.6.8) was adopted to solve the constrained optimization problem. Gurobi solver (version 9.0.2) was used for the calculation of all the linear program (LP) and mixed integer linear program (MILP) problems formulated in this work (Gurobi Optimization and LLC, 2020).

## 3. Results

### 3.1. The influence of different constraints on predicted growth rates

In order to determine a proper *K* value for thermodynamic constraints according to equation (8), we analyzed the Df_i_ variability for all the 1157 reactions with 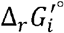 parameters in the irre_iML1515 model, and determined that a *K* value of 1249 kJ/mol is appropriate. At the same time, 24 thermodynamically unfavorable reactions were obtained (Table D, maxDf_i_ < 0). Therefore, the 24 reactions cannot form feasible pathways (Df_i_≥0) predicted by EcoTCM and EcoETM. However, the results of flux variability analysis (FVA) for the pathways with the maximum growth rate predicted by the iML1515 model showed that the two thermodynamically unfavorable reactions E4PD_reverse (catalyzed by erythrose 4-phosphate dehydrogenase) and CBMKr (catalyzed by carbamate kinase) are involved in optimal pathways. Similarly, the two thermodynamically unfavorable reactions DXYLTD_reverse (catalyzed by D-xylonate dehydratase) and CBMKr, are necessary for pathways leading to the maximum growth rate predicted by EcoECM. According to these results, the solution space of iML1515 and EcoECM can be reduced by adding thermodynamic constraints by only excluding individual thermodynamically unfavorable reactions.

**Fig. 1.**
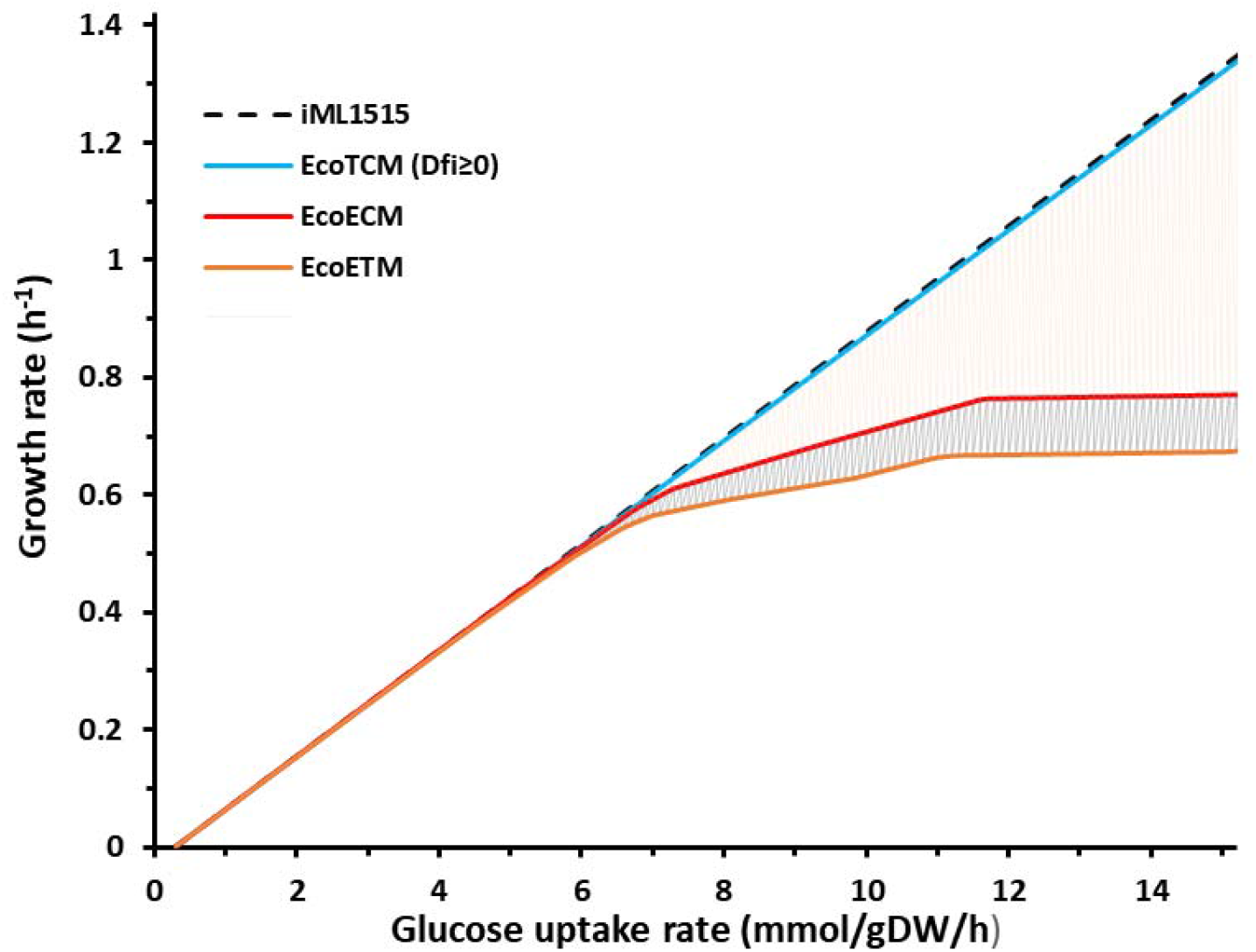
The maximum growth rates predicted by different models. iML1515 model (black dotted line); EcoTCM (blue line); EcoECM (red line) and EcoETM (orange line).

Metabolic network models are often used to predict growth phenotypes and to detect product synthesis pathways (Trudeau et al., 2018; Yang et al., 2019). The integration of enzymatic and thermodynamic constraints into the GEMs is expected to produce more biologically feasible results by reducing the process of subsequent evaluation, screening and verification. Therefore, we further compared the optimal growth calculated based on the iML1515 model and the models integrating these two kinds of constraints separately and simultaneously. As shown in Fig. 1, integrating thermodynamic constraints alone (EcoTCM) does not have any apparent effect on the predicted growth rates, while enzymatic constraints had a more dramatic impact on the predicted growth rates. Furthermore, they also amplify the effect of the thermodynamic constraints as shown by the apparent differences between the results of EcoETM and EcoECM. This indicates that more thermodynamically unfavorable and enzyme costly pathways were excluded from the solution space by integrating both constraints, resulting in more realistic pathway prediction. It should also be noted that the growth rates are mainly constrained by substrate availability at low substrate consumption rates. Therefore, the new constraints mainly affected the calculated optimal growth rates at higher substrate consumption rates.

To verify the thermodynamic feasibility of the pathways from the models, we calculated the MDF of pathways using the optMDFpathway method (Hadicke et al., 2018), which required preset growth rates. We gradually increase the expected rate of growth (by adjusting the lower bound of biomass synthesis reaction fluxes), and then solved the MDF of the pathways before and after adding constraints. In Fig. 2, the black dotted line indicates the maximum growth rate predicted by the iML1515 model when the glucose uptake rate is set at 10 mmol/gDW/h. On this basis, the optMDFpathway method was used to calculate the MDF distribution of biomass synthesis pathways. The results revealed that in the whole feasible space of growth rates (left side of the black dotted line), there is at least one thermodynamically feasible pathway (MDF≥0) that can achieve the optimal thermodynamics (MDF = maxDf_i_ = 2.667 kJ/mol). After introducing enzymatic constraints, the result showed a similar trend that the MDF of pathways decreased gradually with the growth rate, and the feasible space was reduced significantly, indicating that at high growth rates (such as ≥0.63 /h), a certain number of pathways satisfying the enzymatic constraints are not thermodynamically feasible.

**Fig. 2.**
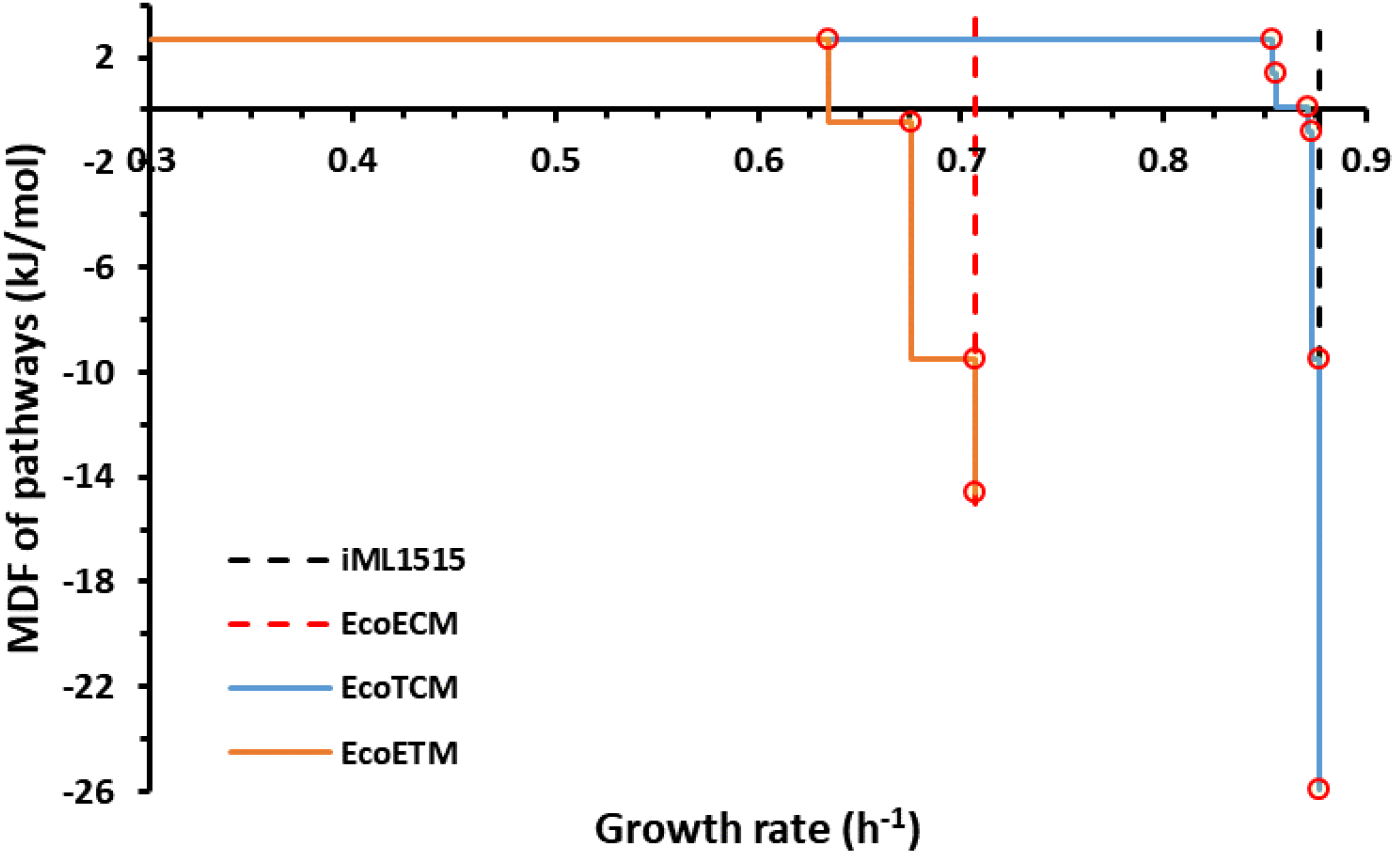
The optimal thermodynamic driving force (MDF) of biomass synthesis pathways under different constraints. The maximum yields predicted by the iML1515 (black dotted line), EcoTCM (blue line), EcoECM (red dotted line) and EcoETM (orange line) models are shown. The points where the MDF suddenly changes are circled in red.

### 3.2. Analysis of bottleneck reactions, limiting metabolites and key enzymes

One application of MDF is to identify the bottleneck reactions and limiting metabolites, which in turn can help propose specific targets for pathway control and optimization (Dash et al., 2019; Yang et al., 2019). On the other hand, ECGEM can predict the optimal enzyme distribution, and thus discover the key enzymes in a pathway as engineering targets (Zheng et al., 2017). As both the bottleneck reactions and key enzymes depend on specific conditions (Trondle et al., 2020), we selected ten turning points (Fig. 2, circled in red) to illustrate the analysis method of bottleneck reactions, limiting metabolites and key enzymes in the ETGEMs framework and to explore the possible reasons for the reduction of solution space by different constraints in detail.

Specifically, we fixed the growth rate at the maximum value that can meet a MDF (B value), and then performed Dfi variability analysis for the reactions constrained by thermodynamics. Hence, we calculated the maxDf_i_ and minDf_i_ of every reaction. When both the growth rate and MDF are preset at maximum values, if the Df_i_ of a reaction does not have variability (ΔDf_i_ = maxDf_i_ – minDf_i_, = 0) and is equal to the B value, it is a bottleneck reaction (Hadicke et al., 2018). Then, the variability of x_i_, which characterizes the metabolite concentration and the variability of enzyme costs, was analyzed in the same way.

In the research of Hadicke et al., the reaction of CBMKr is thermodynamically unfavorable and its stoichiometric relationship is controversial, so the reaction CBPS catalyzed by carbamoyl-phosphate synthase was used to replace the CBMKr as the only way to synthesize Cbp. It should be noted that when CBMKr is allowed to participate in a pathway, the MDF of the pathway should be reduced to below −9.49 kJ/mol (Table D), as mentioned above. Because the enzyme efficiency of CBPS is not considered in EcoTCM, the replacement will not have a particularly significant impact on the growth rate. Similarly, the fact that CBMKr is thermodynamically unfavorable is ignored in EcoECM, so it is allowed to participate in the biomass synthesis process. However, when considering both the thermodynamic and enzymatic constraints in EcoETM, CBMKr was excluded because of its poor thermodynamics, and the problem of low efficiency of CBPS (Guillou et al., 1992) was highlighted simultaneously. As shown in Table E (in Supplementary file2) and Table 2, the CBPS reaction had the highest enzyme cost due to poor kinetic parameters (accounting for 6.9% of the total enzyme cost of the whole pathway), indicating that the replacement of the two reactions actually has a significant impact on growth. It can be seen that the thermodynamic and enzymatic constraints offer two different perspectives on the control steps of a pathway, so the key reactions determined by the two approaches may be very different. Therefore, EcoETM can anchor the thermodynamic bottlenecks and enzymatic key steps of a pathway more effectively, which is conducive to the accurate and comprehensive optimization of a pathway.

**Table 2.** Comparison of parameters for the two Cbp synthesis reactions

| Reaction ID | Reaction Equation | $k_{cat}/MW$ (reverse)<br>$h^{-1} \cdot kDa^{-1}$ | $d_r G'^0$<br>kJ/mol | $Df_i$ range<br>kJ/mol |
| --- | --- | --- | --- | --- |
| CBPS | $2.0 \text{ atp\_c} + \text{gln\_L\_c} + \text{h2o\_c} + \text{hco3\_c} \rightarrow 2.0 \text{ adp\_c} + \text{cbp\_c} + \text{glu\_L\_c} + 2.0 \text{ h\_c} + \text{pi\_c}$ | 54.13 | $-13.4 \pm 5.2$ | $-21.65 \sim 105.48$ |
| CBMKr | $\text{atp\_c} + \text{co2\_c} + \text{nh4\_c} \rightleftharpoons \text{adp\_c} + \text{cbp\_c} + 2.0 \text{ h\_c}$ | 4179.36<br>(1221.59) | $19 \pm 7.8$ | $-81.96 \sim -9.49$ |

In addition, the reaction TPI is also a crucial reaction from both the thermodynamic and enzymatic perspectives (Tables S2 and S4, in Supplementary file1). The higher enzyme usage caused by high flux indicates its importance in the biomass synthesis process. At the same time, it is a reversible reaction that is prone to reaching an equilibrium state, and its product glyceraldehyde 3-phosphate (g3p) is also the substrate of another bottleneck reaction, GAPD (Table F, in Supplementary file2). Therefore, the concentration of g3p is strictly trapped. There is a close relationship between bottleneck reactions and limiting metabolites, and the sharing of metabolites among reactions is an important reason for the phenomenon of distributed bottleneck reactions (Hadicke et al., 2018; Mavrovouniotis, 1993), which suggests that we need to weigh the potential and comprehensive impact of bottleneck reactions when developing an optimization strategy according to an optimal distribution of metabolite concentrations.

Pathway evaluation performed in previous studies (Trudeau et al., 2018; Yang et al., 2019) indicated that although the theoretical maximum yield remains unchanged, many pathways should still be excluded due to criteria related to enzyme kinetics and thermodynamics. In section 3.1, the effect of thermodynamic constraints on reducing the yield space was not always apparent in Fig. 1. Because the number of solutions is more representative of the size of solution space than the maximum yield, the results do not necessarily indicate that thermodynamic constraints play a dispensable role in reducing the solution space. Based on this analysis, we can see that thermodynamic constraints can screen feasible pathways by either 1) determining the thermodynamically unfavorable reactions, thereby excluding all the pathways in which they are necessary, such as CBMKr, DXYLTD_reverse and E4PD_reverse, or 2) by eliminating reactions that are thermodynamically feasible in principle, but no longer meet the criteria due to shared limiting metabolites, which lead to distributed bottleneck reactions. For example, the simultaneous occurrence of PGCD, GAPD, FBA, PGK and TPI precludes the feasibility of biomass synthesis (Tables E-G in Supplementary file2).

### 3.3. Analysis of products synthesis pathways using the four models

To further investigate the phenotypic prediction differences between models with different constraints, we reanalyzed the pathways for the synthesis of 20 products with the highest yield from glucose used in another study (Hadicke et al., 2018). The prediction results of the iML1515 model show that Cbp is the product with the highest yield. Since Cbp is an essential precursor of L-arginine (L-Arg), we also calculated the yield of L-Arg and its other precursor, ornithine (Orn). As shown in Fig. 3A, the calculated optimal product synthesis rates from iML1515 for the 22 products all had linear relationships with the glucose uptake rate, and the synthesis rate of Cbp was much higher than those of other products. As shown in Fig. 3B, with the addition of thermodynamic constraints, the rate still increased linearly, but the synthesis rates for Cbp, Orot, Dhor_S, Cbasp, Orot5p and L-Arg were lower than in iML1515 at the same glucose uptake rates (also shown for individual products in Fig. 4). By analyzing the MDF change curves of these products, we found that all the MDF values of the maximum yield pathways predicted by iML1515 were −9.49 kJ/mol (Figure B, in Supplementary file2) due to the participation of the bottleneck reaction CBMKr. This very low value indicated that CBMKr is thermodynamically unfavorable, and it was automatically excluded from the model with thermodynamic constraints, generating more realistic pathway predictions than the iML1515 model for Cbp-derived products such as Orot, Dhor_S, Cbasp, Orot5p and L-Arg.

**Fig. 3.**
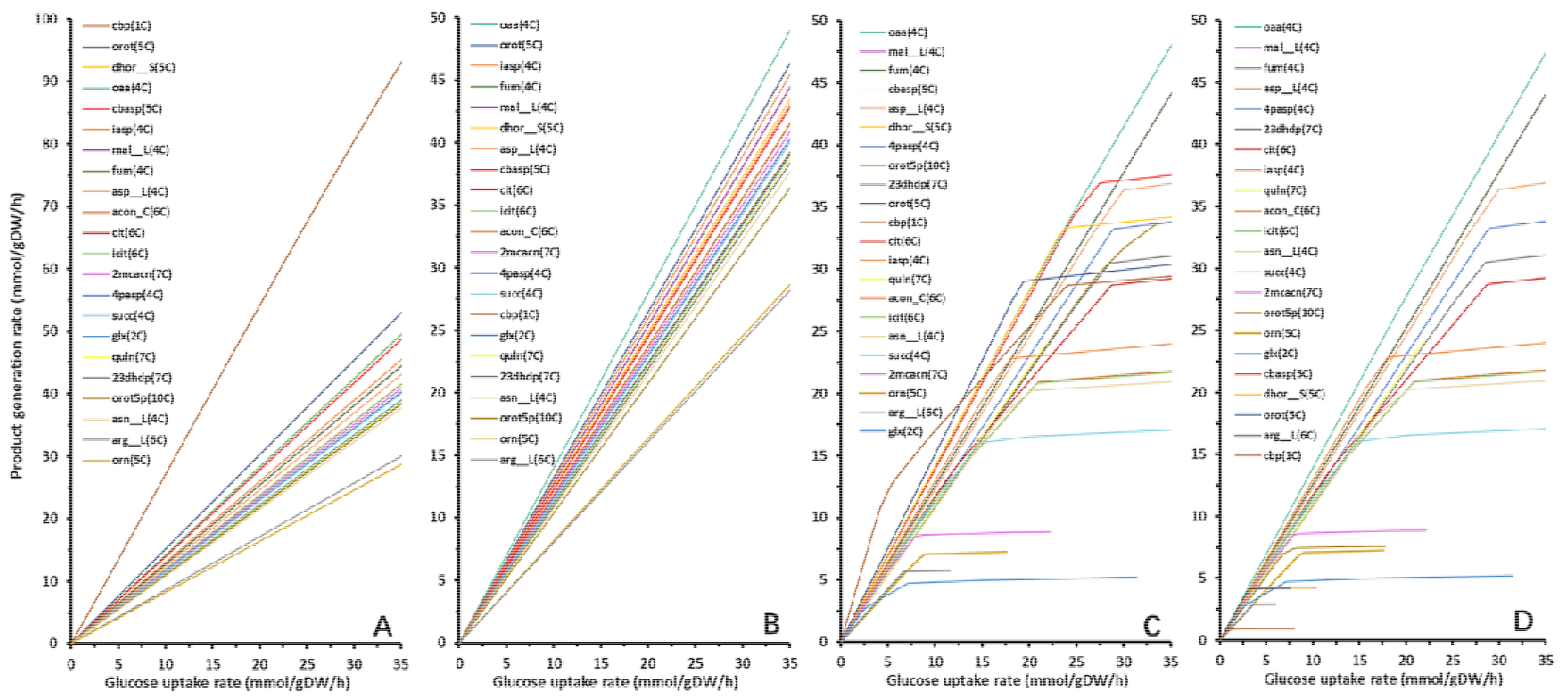
The simulation results of 22 product synthesis rates based on various models. **(A)** iML1515; **(B)** EcoTCM; **(C)** EcoECM; **(D)** EcoETM. The order of names in the legend is the same as the order of the final values of the production curves (from top to bottom). The molar amount of products was normalized based on glucose (6 C-atoms).

**Fig. 4.**
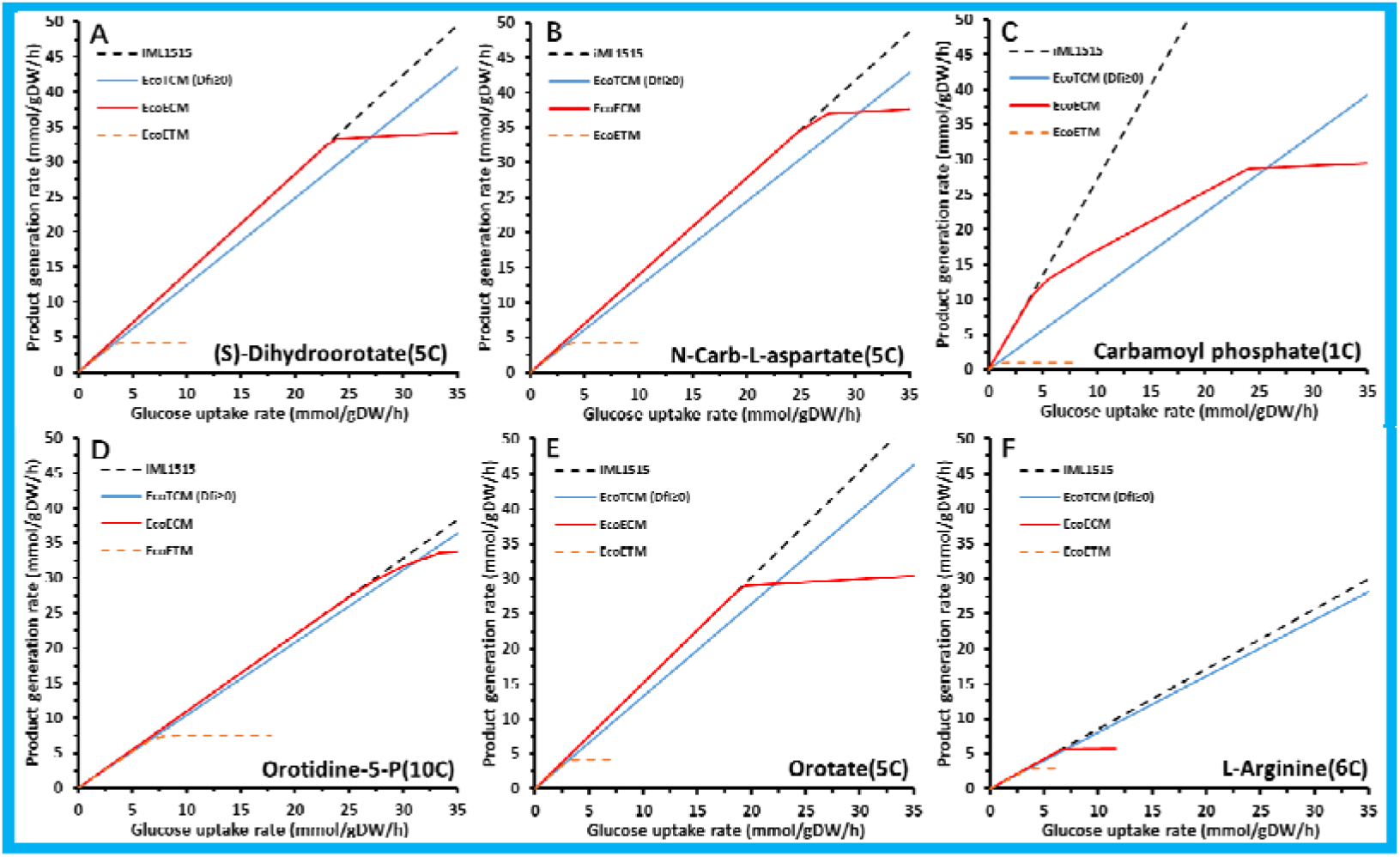
The predicted synthesis rates for various products by various models. **(A)** (S)-Dihydroorotate (Dhor_S); **(B)** N-Carb-L-aspartate (Cbasp); **(C)** Carbamoyl phosphate (Cbp); **(D)** Orotidine-5-P (Orot5p); **(E)** Orotate (Orot); **(F)** L-Arginine (L-Arg). The molar amount of products was normalized based on glucose (6 C-atoms).

In addition to these Cbp-derived products, the calculated maximum rate of oxaloacetate (Oaa) also decreased slightly (Figure A and Table H, in Supplementary file2) in the models with thermodynamic constraints. After setting the Oaa synthesis rate at the maximum value predicted by iML1515 and solving the MDF of the Oaa synthesis pathway(s), an MDF of −0.632 kJ/mol was obtained (Figure B, in Supplementary file2). Then, by analyzing the Df_i_ variability, the three reactions FLDR2 (catalyzed by flavodoxin reductase), PFL (catalyzed by pyruvate formate lyase) and POR5_reverse (catalyzed by pyruvate synthase) were identified as distributed bottlenecks (Mavrovouniotis, 1993). Due to the shared metabolites between the bottleneck reactions, the simultaneous participation of the three reactions would preclude the thermodynamic feasibility of Oaa biosynthesis (Df_i_<0).

At low substrate uptake rates, the predicted synthesis rates are limited by substrate availability and there is no difference in the rates from iML1515 (Fig. 3A) and those from the enzyme constrained model (Fig. 3C). When the substrate uptake rate was increased, the rate curves in Fig. 3C began to turn, indicating that the enzyme availability starts to be a limiting factor, and the pathways with lower enzyme costs need to be enabled to satisfy the enzymatic constraints. The minimum enzyme cost of the optimal Cbp synthesis pathway calculated based on iML1515 was 65.63 mg /(mmol glucose/h). With a total enzyme constraint of 0.228 g enzyme/gDW, the maximum rate of glucose uptake of this pathway was calculated to be 3.47 mmol/gDW/h. Therefore, at glucose uptake rates above this value, this high enzyme cost pathway is gradually switched to new pathways with lower enzyme costs and lower yields (Fig. 3C). When the glucose uptake rate was set to 1 mmol/gDW/h, the Cbp synthesis flux was set to 15.67 mmol/gDW/h, and the total enzyme amount was set to 133.25 mg/gDW, the minimum enzyme cost of reactions in the pathway showed that AKGDH (catalyzed by 2-oxogluterate dehydrogenase), GLCptspp (realized by the PTS system) and CBMKr were the three reactions with the highest enzyme cost, at 8.10, 5.50 and 3.75 mg/gDW, respectively. The main reasons for the high enzyme cost were the high protein molecular weight (AKGDH, 2418.39 kDa), low *k*_cat_ value (GLCptspp, 10.6 /s) and high flux demand (CBMKr, 15.67 mmol/mmol glucose), respectively.

As shown in Fig. 3D, due to the integration of both thermodynamic and enzymatic constraints, the synthesis rate of some products decreased significantly. Cbp decreased from the highest rate predicted by the iML1515 model to the lowest one predicted by EcoETM, showing the combined effect of the two constraints on the feasibility of pathways and the great reduction of the solution space. By comparing Figs. 3A-D, it can be seen that the integration of thermodynamic constraints (Fig. 3B) and enzymatic constraints (Fig. 3C) did affect the prediction results of iML1515 (Fig. 3A) from different perspectives. At the initial stage of glucose uptake, thermodynamic constraints can change the yield and ranking order of product synthesis. When the glucose uptake rate reaches a specific level, the enzyme amount becomes the limiting factor, and the rate curves from EcoECM begin to show differences with those form the iML1515 model. After integrating the two constraints in one model, many unfavorable pathways were excluded from EcoETM, which led to a much smaller solution space and more precise prediction of pathways. For example, the maximal arginine synthesis rate of the thermodynamically feasible and low enzyme cost pathways is actually quite low (is 4.36 mmol/gDW/h at the maximum glucose uptake rate of 6 mmol/gDW/h) and significantly different from the results predicted by iML1515 (the carbon yield reduced by 43.1%, Fig. 4F).

The MDF change in Fig. 5 clearly shows the switching of pathways under thermodynamic constraints. Due to the participation of CBMKr, the MDF suddenly drops in the synthesis process of Cbp (Fig. 5A), as well as its derivatives, such as L-arginine (Fig. 5B). As can be seen in Fig. 5C, although there is no thermodynamically unfavorable reaction in the Oaa synthesis pathway, the simultaneous occurrence of 3 distributed bottleneck reactions, FLDR2, PFL and POR5_reverse, nevertheless precludes the thermodynamic feasibility of the pathway. It should be noted that by adjusting the threshold of MDF, i.e. by introducing a strong thermodynamic constraint such as increasing the MDF threshold from 0 to 1 kJ/mol (Trudeau et al., 2018), more pathways can be excluded from the solution space, allowing the prediction of more thermodynamically feasible pathways(Fig. S1, in Supplementary file1). In addition, by comparing Figs. 5D and E, it can be seen that iML1515 and EcoTCM cannot distinguish the synthesis curves of citrate (Cit) and isocitrate (Icit), while EcoECM (including EcoETM) can distinguish them. The synthesis of Icit from Cit requires an additional reaction catalyzed by aconitase (consisting of the two half-reactions ACONTa and ACONTb in the model), so more enzyme is needed for the Icit synthesis process, leading to earlier pathway switching than for Cit. However, the two reactions are not thermodynamic barriers and there is no carbon and energy loss, so their production curves in EcoTCM and iML1515 are identical.

**Fig. 5.**
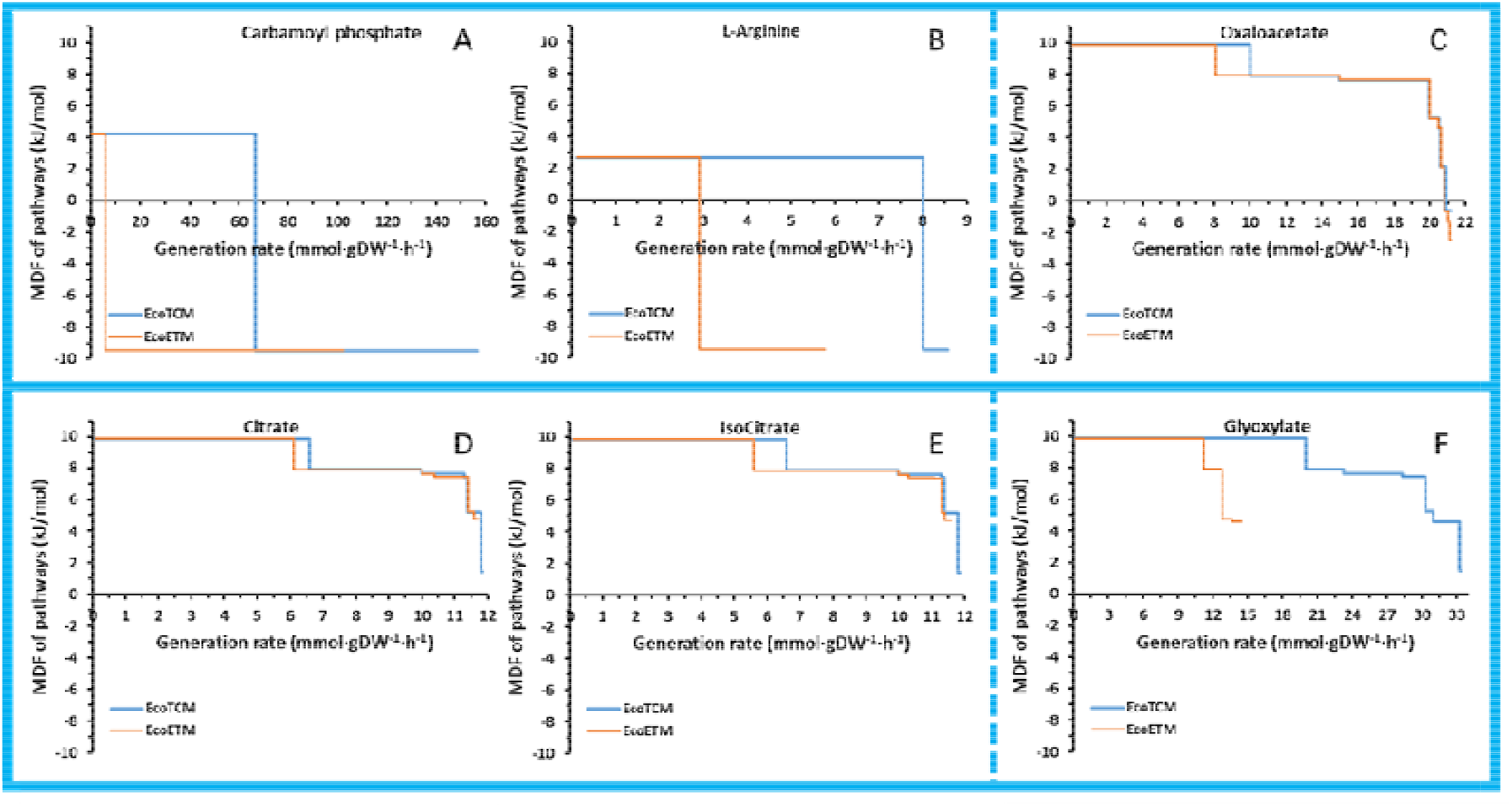
The MDF of product synthesis pathways under different constraints. **(A)** Carbamoyl phosphate; **(B)** L-Arginine; **(C)** Oxaloacetate; **(D)** Citrate; **(E)** Isocitrate; **(F)** Glyoxylate.

To further investigate the differences of the predicted optimal pathways from iML1515 and EcoETM, we plotted the calculated L-Arg synthesis pathways from the two models with a glucose uptake rate of 3.64 mmol/gDW/h (the turning point at which the enzyme constraint becomes the limiting factor), as shown in Fig. 6. The L-Arg production rates at this point were 3.12 mmol/gDW/h based on iML1515 and 2.90 mmol/gDW/h based on EcoETM. As can be seen in Fig. 6, the key difference is in the Cbp production part. In the pathway obtained from iML1515, the CBMKr reaction with high enzyme efficiency is used (Fig. 6A). By contrast, this reaction is not in the pathway from EcoETM because it is thermodynamically unfavorable, and the CBPS reaction with low carbon yield and low enzyme efficiency is used instead (Fig. 6B). This inevitably leads to high enzyme cost of the pathway and a small maximal production rate, which was significantly lower than that predicted by iML1515. This enzyme was therefore identified as an engineering target for improving arginine production. In addition, through the Df_i_ variability analysis, we also found a thermodynamic bottleneck reaction in the L-Arg synthesis pathway, catalyzed by acetylglutamate kinase (ACGK, its maxDf_i_ is only 2.667 kJ/mol). Furthermore, ACGK is also the thermodynamic bottleneck for biomass synthesis, as described in 3.2 and 3.3. Its thermodynamic feasibility is likely to be highly dependent on ADP depletion reactions, such as pyruvate kinase (Vogel and McLellan, 1970). Coupling between reactions is an important means to overcome the thermodynamic bottleneck for the engineering practice (Zhang et al., 2017)

**Fig. 6.**
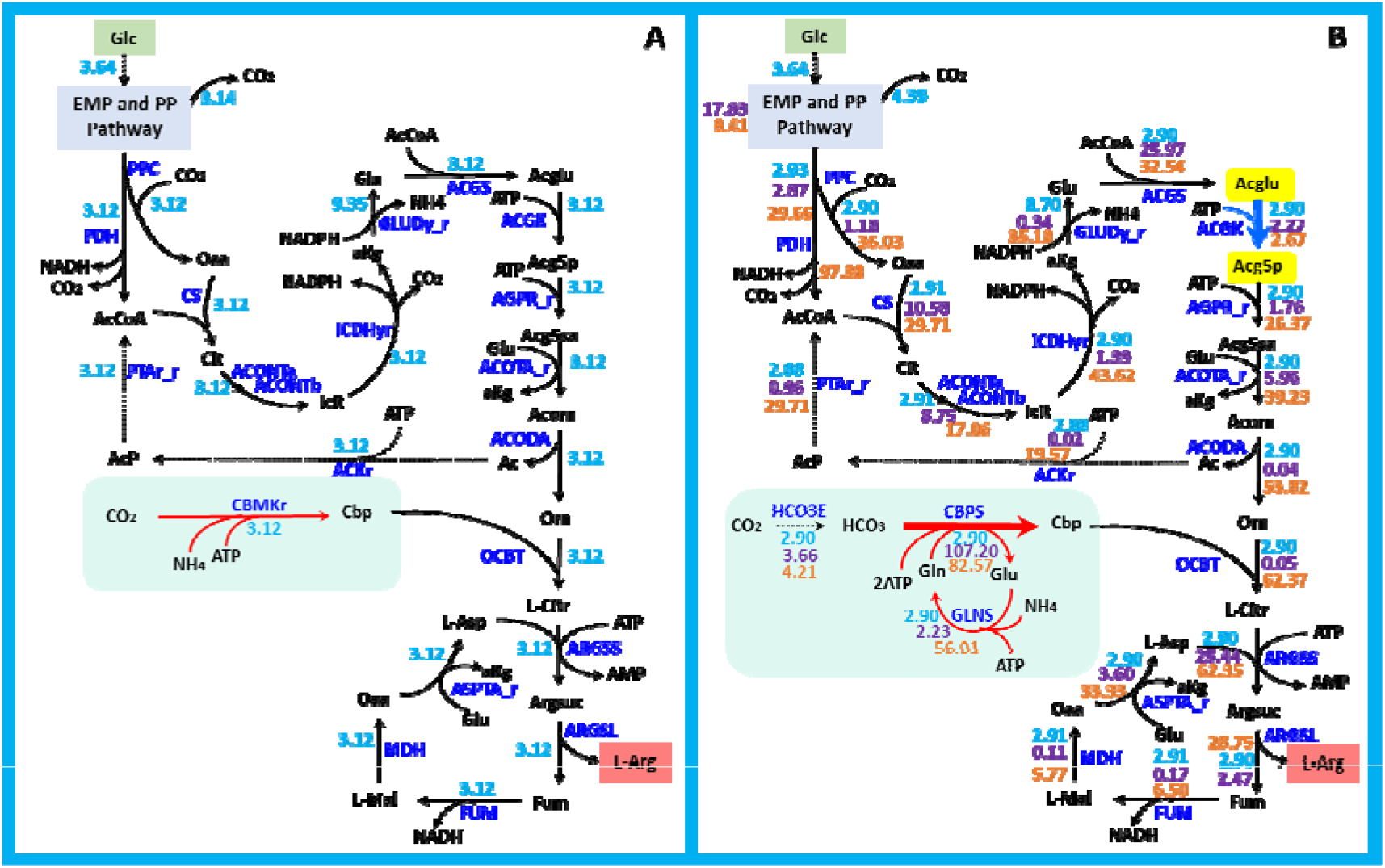
Prediction of the L-arginine synthesis pathway by iML1515 (A) and EcoETM (B). Shown are: the structural change of the pathway (light blue region); the reaction with the highest enzyme cost (red thick arrow); the thermodynamic bottleneck reaction (blue arrow); the limiting metabolite (yellow background); and the simplified pathways of EMP and PP (navy-blue background). The unit of the flux value is mmol/gDW/h (blue, on the top); the unit of the enzyme cost is mg/(mmol glucose /h) (purple, in the middle); and the unit of the maximum thermodynamic driving force is in kJ/mol (orange, at the bottom).

## 4. Discussion

The development of ETGEMs benefits from the excellent biological basis and mathematical modeling foundation of ECGEMs (Adadi et al., 2012; Sanchez et al., 2017) and thermodynamic constraint models (Hadicke et al., 2018; Henry et al., 2007; Salvy et al., 2019). In ETGEMs, enzyme restriction leads to a decrease of the predicted maximum yield by excluding pathways with high enzyme costs. Accordingly, the cells have to switch to new pathways to satisfy the enzymatic constraint (Chen and Nielsen, 2019). The addition of thermodynamic constraints can not only limit the feasibility of pathways, but also optimize the thermodynamic feasibility of bottleneck reactions in the pathway by adjusting the concentration of metabolites, and predict the MDF for a pathway. With the addition of thermodynamic and enzyme constraints, ETGEMs strengthen the restriction of the feasibility of a pathway to allow more realistic pathway prediction. It can also be used to identify thermodynamic bottleneck reactions and low efficiency enzymes, and thus provide guidance for pathway engineering.

Computational methods have been used to systematically design novel pathways in recent studies. It is often necessary to screen pathways based on certain criteria to choose the most promising pathways for experimental verification (Trudeau et al., 2018; Yang et al., 2019). Pathway evaluation needs to integrate thermodynamic and kinetic standards directly in the GEMs. Therefore, by integrating the dual constraints into the GEMs, the thermodynamic and enzymatic cost of the pathway can be calculated. Taking the L-Arg synthesis pathway as an example, in addition to flux distribution, thermodynamic bottleneck reactions, limiting metabolites, enzyme cost distribution and key enzyme information are also given. Therefore, the ETGEMs, if combined with certain algorithms, are expected to be an effective tool for systematic pathway design. In addition, the integration of thermodynamic constraints in the reaction sets of a specified model can avoid the repetitive preparation of pathway information.

The values of parameters such as 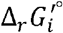 and *kcat* can greatly affect the prediction results of a constrained model. In the construction of EcoETM, some standard thermodynamic parameters were not successfully estimated due to the inconsistent names of metabolites or the lack of KEGG reaction IDs in the iML1515 model. Besides, in order to improve the coverage of the 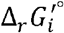 parameters, some approximate 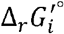 values were obtained by neglecting the groups in the metabolites that have not changed and do not have the evident role of thermodynamic promotion (e.g. GPDDA2: replacing “Glycero-3-phosphoethanolamine + H2O <=> Ethanolamine + Glycerol 3-phosphate” with “Ethanolamine phosphate + H2O <=> Ethanolamine + Orthophosphate”), by referring to similar reactions with the same group changes (e.g. GP4GH: replacing “GppppG + H2O <=> 2 GDP” with “AppppA + H2O <=> 2 ADP”), or by replacing the metabolites that cannot be evaluated with structural similarly metabolites (e.g. L_LACD3: replacing “Menaquinone 8” with “Menaquinone”). This is a preliminary exploration of the possibility of parameter reference between the reactions due to the similarity of the involved compounds and changed groups. In fact, more accurate larger-scale enhancement of 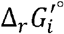 parameter coverage will still depend on the resources of reactions and parameters available in databases such as eQuilibrator, TECRDB (Goldberg et al., 2004) and KEGG, as well as the combination of efficient methods, such as machine learning (Heckmann et al., 2018). Researchers have made efforts to calibrate parameters by referring to the yield and flux distribution of the biomass and product synthesis processes (Bekiaris and Klamt, 2020), or reasonably narrowing metabolite concentration ranges according to metabolomic data (He et al., 2020). The improvement of parameter accuracy and coverage will increase the prediction efficiency, reduce the cost of result evaluation, and contribute to the construction of powerful metabolic models of *E. coli* and other species.

## 5. Conclusions

In this work, we developed a novel functional modeling framework for genome-scale metabolic models with integrated enzymatic and thermodynamic constraints, named ETGEMs. The pathway calculation results indicated that many thermodynamically unfavorable and enzymatically costly pathways were excluded by the new constraints, leading to more realistic pathway prediction. By comparing the pathways from different models, the thermodynamic and enzymatic bottlenecks in the pathways can be identified, providing new targets for directed evolution and metabolic engineering.

## Acknowledgments

This work was funded by the National Key Research and Development Program of China (2018YFA0900300, 2018YFA0901400); the International Partnership Program of Chinese Academy of Sciences (153D31KYSB20170121). We thank Elad Noor, Steffen Klamt, Axel von Kamp for providing more details about their work. We thank David Heckmann for sharing EC number file with us. We thank Dr. Chaoyou Xue in revising the manuscript.

